# Decreased biofilm formation in *Proteus mirabilis* after short-term exposure to a simulated microgravity environment

**DOI:** 10.1101/2021.04.07.438892

**Authors:** Dapeng Wang, Po Bai, Bin Zhang, Xiaolei Su, Xuege Jiang, Tingzheng Fang, Junfeng Wang, Changting Liu

**Author notes:** Corresponding authors: Changting Liu, Respiratory Diseases Department, The Second Medical Center of PLA General Hospital, Beijing, 100853, China, Email address, Junfeng Wang, Respiratory Diseases Department, The Second Medical Center of PLA General Hospital, Beijing, 100853, China. These authors contributed equally to this work (Dapeng Wang, Po Bai, Bin Zhang).

## Abstract

**Background:** Microbes threaten human health in space exploration. Studies have shown that *P. mirabilis* has been found in human space habitats. In addition, the biological characteristics of *P. mirabilis* in space have been studied unconditionally. The simulated microgravity environment provides a platform for understanding the changes in the biological characteristics of *P. mirabilis*.

**Objective:** This study intends to explore the effect of simulated microgravity on *P. mirabilis*, the formation of *P. mirabilis* biofilm and its related mechanism.

**Methods:** The strange deformable rods were cultured continuously for 14 days under the microgravity simulated by (HARVs) in a high-aspect ratio vessels. The morphology, growth rate, metabolism and biofilm formation of the strain were measured, and the phenotypic changes of *P. mirabilis* were evaluated. Transcriptome sequencing was used to detect differentially expressed genes under simulated microgravity and compared with phenotype.

**Results:** The growth rate, metabolic ability and biofilm forming ability of *P. mirabilis* were lower than those of normal gravity culture under the condition of simulated microgravity. Further analysis showed that the decrease of growth rate, metabolic ability and biofilm forming ability may be caused by the down-regulation of related genes (*pstS,sodB* and *fumC*).

**Conclusion:** It provides a certain reference for the prevention and treatment of *P. mirabilis* infection in the future space station by exploring the effect of simulated microgravity exposure on *P. mirabilis*.

## 1. Introduction

Manned space technology is developing rapidly. Astronauts and cabin equipment inevitably bring microbes, including bacteria, fungi and viruses, into space in the process of manned spaceflight. The space environment is characterized by a series of prominent features more than 100km above the earth’s surface, including strong ionizing radiation, high vacuum, ultra-low temperature, ultra-high temperature and weightlessness (Thirsk et al., 2009, Gerald et al., 2009, Sas et al., 2005, Song et al., 2005). Various factors in the space environment can induce genetic changes of microorganisms, and then affect the phenotypic characteristics of microorganisms, including morphology, growth rate, biofilm forming ability, virulence, drug resistance and so on, which may induce infectious diseases and affect the health of resident personnel. As a consequence, it is of great significance to study the phenotypic characteristics of microorganisms in space environment. *P. mirabilis* belongs to Proteus and widely exists in soil, water and feces (Branka et al., 2016). In addition, *P. mirabilis* is highly pathogenic and is one of the main causes of persistent and refractory urinary tract infections. Studies have shown that *P. mirabilis* has been tested for in space and post-flight astronaut samples in human habitats (Chaffer et al. 2011, Castro et al., 2004). The occurrence of infectious diseases is more likely to occur due to the decline of human immunity in the space environment. As a consequence, how to prevent and treat the infection caused by *P. mirabilis* in the space environment is worthy of attention. However, studies on the biological characteristics of *P. mirabilis* in the space environment have not been carried out due to concerns about the cost, time and safety of spacecraft flight.

Microgravity environment refers to the environment in which the apparent weight of the system is much smaller than its actual weight under the action of gravity. In fact, it is the most important environmental factor encountered in the process of manned space flight. Microgravity simulation experiment has become an important and necessary preparation for the development of manned spaceflight. The biological characteristics of microorganisms under simulated microgravity have been widely studied in recent years. Some results show that under the simulated microgravity conditions, microorganisms show the same biological characteristics as the space environment. For example, it was found that the virulence of *Salmonella typhimurium* increased under simulated microgravity, which was consistent with that of *Salmonella typhimurium* under space microgravity. Further studies at the molecular level also show that RNA-binding protein Hfq is involved in the process of simulating the effects of microgravity and space environment. Simulated microgravimeter was used to observe the changes of biological characteristics and internal mechanism of *P. mirabilis* under microgravity. It can provide a strategy for the prevention and treatment of infectious diseases caused by *P. mirabilis* in space environment in the future.

## 2. Materials and Methods

### 2.1 Bacterial strains and culture conditions

The whole study used *P. mirabilis* clinically separated from the urine of a male patient. The API20NE identification system (BioMèrieux, Craponne, France) and 16S rDNA gene sequencing were applied to identify the strains. The *P. mirabilis* strains grew aerobically in Luria-Bertani (LB) medium (with 0.5% agar) at 37 °C. The experimental (simulated microgravity, SMG) and the normal gravity control (NG) were set up in a high-aspect rotating vessel (HARV) bioreactor (Synthecon, Inc., Houston, Texas, USA) and cultured continuously. Figure 1 shows SMG culturing by rotating the bioreactor with the axis perpendicular to gravity, while the cultivation under NG is realized in case of its axis parallel to gravity (Nickerson et al., 2000). *P. mirabilis* was inoculated in the HARV bioreactor with a dilution of 1: 200. Each bioreactor was fully filled with fresh LB medium of ∼55 ml, and the bubbles were removed carefully. After culturing in HARVS at 37 °C for 24 h at 25 rpm, the P. mirabilis cultures under SMG and NG were diluted and transferred into another HARVS fully filled with LB medium. *P. mirabilis* inoculation was cultured in the HARV bioreactors for 2 weeks. *P. mirabilis* cultured under SMG was called PMML strain, while that cultured under NG was called PMGL strain. The PMML and PMGL groups were serially diluted in phosphate-buffered saline (PBS), plated on LB agar, and counted. The cultures obtained were tested in various ways.

**Figure 1.**
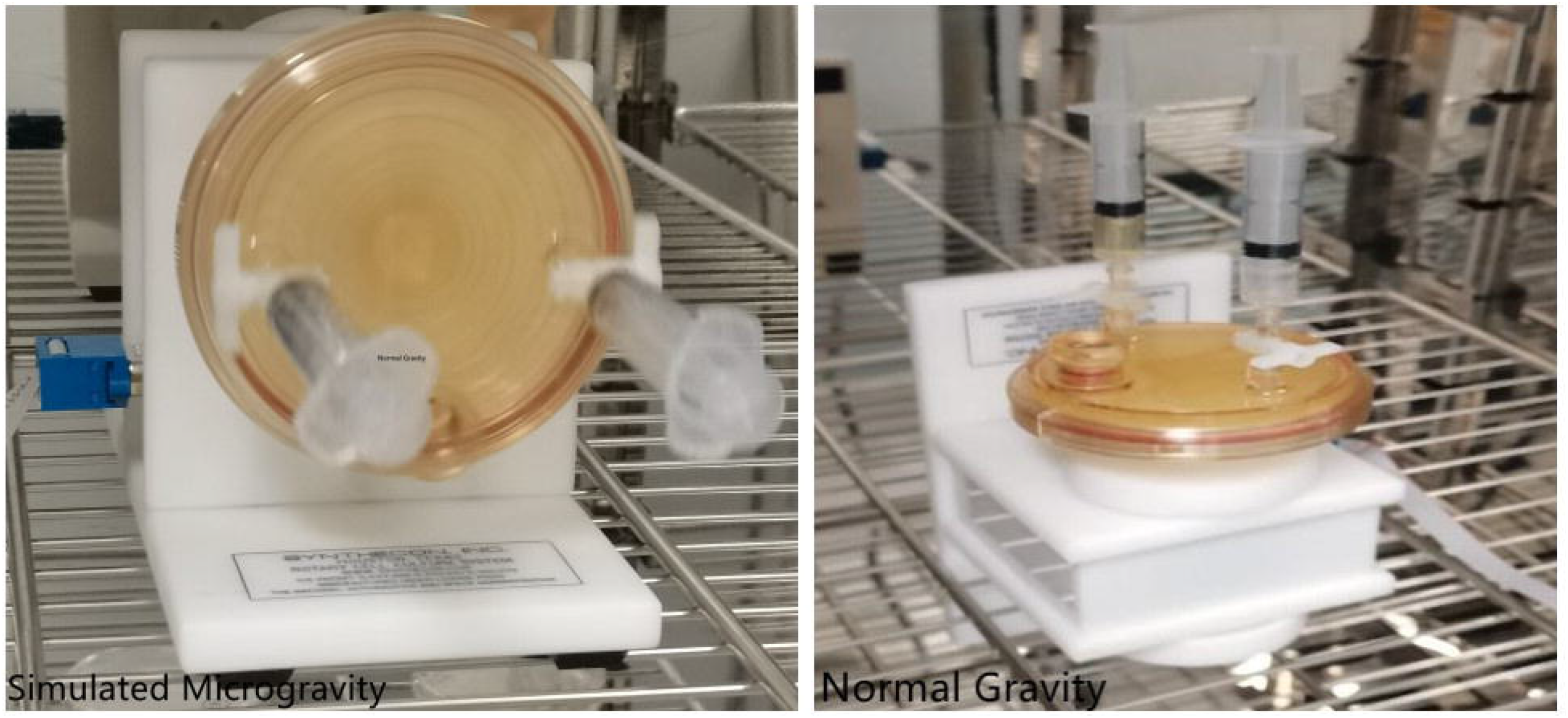
Experimental setup of the high-aspect rotating vessels bioreactors. *P. mirabilis* cells in the HARV bioreactor are grown under the simulated microgravity (SMG) condition with its axis of rotation perpendicular to gravity or grown under the normal gravity (NG) with its axis of rotation vertical to gravity. The bioreactors are filled with LB medium and the bubbles are removed.

### 2.2 Phenotypic analysis

#### 2.2.1 Scanning electron microscopy (SEM)

PMML and PMGL strains were cultured in LB medium, washed with sterile phosphate-buffered saline (PBS; pH 7.4), and fixed with 4% glutaraldehyde overnight. The samples were washed again with PBS, dehydrated in increasing grades of ethanol, and then critical pointdried. After coating the specimens with gold-palladium, they were observed using an FEI Quanta 200 scanning electron-microscope (USA).

#### 2.2.2 Growth rate assay

The Bioscreen C system and BioLink software (Lab Systems) were applied to monitor *P. mirabilis* growth. The overnight growing of *P. mirabilis* strains was made in LB medium at 37 °C. Twenty microliters of the overnight culture at a concentration of 10^6^ CFU/ml was cultivated in Bioscreen C 96-well microtiter plates (Lab Systems). The plates containing fresh LB growth medium of 350 µl per well were incubated under continuous shaking at the maximum amplitude for 24 h at 37 °C. The OD_630_ was measured every 2 h. *P. mirabilis* growth experiments were performed in triplicates.

#### 2.2.3 Carbon source utilization and chemical sensitivity assay

The biochemical features of two strains were tested by using the Biolog GENIII MicroPlate (Biolog, CA) and 94 phenotypic tests, including utilization of 71 carbon sources and assays of 23 chemical sensitivity. The overnight growing of *P. mirabilis* strains on agar plates was made at 37 °C. The centrifugal tube containing two strains was washed with PBS and inoculated into the IF-A inoculum (Biolog, CA). A turbidimeter was used to adjust the concentration of *P. mirabilis* suspension to 10^8^ CFU/ml, and the suspension of 100 μl was inoculated into each well of the 96-well plate. After 24-hour incubation at 37 °C, an automated Biolog microplate reader at 630 nm was used to measure the absorbance. The experiment was carried out in triplicates.

#### 2.2.4 Biofilm assay

##### 2.2.4.1 Crystal violet staining (a)

After being diluted (1:100) in 5 ml LB medium, the PMML and PMGL cultures were moved to glass tubes and incubated at 37 °C for 24 h at 200 rpm. The planktonic bacteria were removed. Soon afterwards, deionized water was used to wash each tube for three or four times. Then, 0.1% of crystal violet dye was used to stain the glass tubes for 15 min at 37 °C. The experiment was carried out in triplicates.

##### 2.2.4.2 Crystal violet staining (b)

After adding approximately 200 μl of the *P. mirabilis* suspension culture (1–5 × 10^7^ CFU/ml) to 96-well microculture plates, the mixture was incubated at 37 °C for 24 h. PBS was used to wash the wells for two times. 200 μl of 0.1% crystal violet was employed to stain the cells (Sigma, St. Louis, MO, USA) for 30 min and washed with PBS. Next, the microculture plates were dried, and the dissolution of the stained biomass samples in 95% ethanol was made. The Thermo Multiskan Ascent Instrument (Thermo, USA) was applied to determine OD*570* for each well. The experiments were carried out in triplicates.

##### 2.2.4.3 Analysis of biofilm formation ability via confocal laser-scanning microscopy (CLSM)

The two strains were cultured on LB medium overnight and inoculated in 35-mm confocal dishes (Solarbio, Beijing, China). The plates were incubated at 37°C for 24 hr and stained with a Filmtracer LIVE/DEAD Biofilm Viability Kit (Invitrogen, Carlsbad, CA, USA) in accordance with the manufacturer’s instructions. Three image stacks were obtained from each sample randomly by CLSM using a Leica TCS SP8 microscope with a Lecia TCS SP8 CSU imaging system. All the experiments were carried out in triplicates.

#### 2.2.5 Transcriptomic sequencing and comparison

##### 2.2.5.1 Sequencing and filtering

*P. mirabilis* cells were collected after five-minute centrifugation at 8,000 × g at 4 °C. The RNeasy Protect Bacteria Mini Kit (Qiagen, Germany) was used to extract total RNA from the two strains according to the manufacturer’ s instructions. The 10-minute centrifugation of P. mirabilis was made at 10,000 × g at 4 °C. Chloroform added with the supernatant was mixed for 15 s. After being transferred into a tube filled with isopropanol, the upper aqueous phase was centrifuged at 13,600 × g for 20 min at 4 °C. After removing the supernatant, three -minute centrifugation of the mixture of precipitate and ethanol was made at 12,000 × g and 4 °C. Soon afterwards, the supernatant was discarded, and the 20-second centrifugation of sample was made at 12,000 × g and 4 °C. After removing the residual liquid by air -drying, the dissolution of RNA pellet in RNase-free water was made. The purity of the samples is tested by NanoDropTM. Divalent cations were used to fragment the purified mRNAs into small pieces (∼200 bp). The first-strand of cDNA using reverse transcriptase and random primers was generated by applying RNA fragments. Following this, DNA polymerase I and RNase H were used to create the second-strand of cDNA. The enrichment and quantification of cDNA fragments were made by using PCR amplification and Qubit 2.0 respectively. Lastly, the BGISEQ-500 was used to construct and sequence cDNA libraries. The SOAPnuke software (version 1.5.2) was applied to filter raw reads, and adapter reads and poly N reads were removed from the raw data to acquire clean reads. Clean reads were located to the reference genome by Bowtie2-2.2.3 (Langmead et al., 2012). Gene expression was quantified by HTSeq v0.6.1, and the calculation of the fragments per kilobase of transcript per million mapped reads (FPKM) of each gene was made according to the gene length. Lastly, the mapping of read counts to the genes was made. The DESeq R package (1.18.0) was employed to make differential gene expression analysis of the two strains. Through DESeq analysis, it is considered that genes yielding a p-value of < 0.05 are differentially expressed. The KEGGseq R package was applied to analyze Kyoto Encyclopedia of Genes and Genomes (KEGG) enrichment of differentially expressed genes (DEGs). It was considered that DEGs with a p-value of < 0.05 is statistically significant. KOBAS software was used to assess the statistical enrichment of DEGs in KEGG pathways.

##### 2.2.5.2 Statistical analysis of gene expression values and DEGs

After mapping the clean reads to the reference strain by applying HISAT (version 2.0.4) and Bowtie2 (version 2.2.5) (Li and Dewey, 2011), the gene expression level with RSEM (Langmead et al., 2012) was calculated. Cluster software (version 3.0) was applied to make cluster analysis of gene expression, and Java TreeView (3.0) (Saldanha, et al., 2004) was used to visualize the results of cluster analysis.

##### 2.2.5.3 Functional annotation and enrichment analysis

The genes from the reference sequences in the KEGG database were selected using the BLAST software to compare and annotate gene function. The gene expression profiles of different sample groups were analyzed using cluster analysis. The transcriptome data was analyzed by identifying DEGs according to the KEGG pathway. Gene enrichment analysis was performed using KEGG functional annotation. A *p*-value ≤ 0.05 indicated significant enrichment of DEGs.

#### 2.2.6 Quantitative real-time PCR (qPCR)

Total RNA and random hexamer primers with SuperscriptIII reverse transcriptase (Invitrogen) using qPCR were employed to synthesize cDNA. The StepOnePLUS PCR system (Thermo, USA) with cDNA as the template were used to carry out the experiment in duplicates for each RNA sample. The related fold change of the target genes in the test and controlled RNA samples was determined. The 16S rRNA gene was regarded as an internal reference. The primer sequences used in this study are shown in Table 1.

**Table 1.**
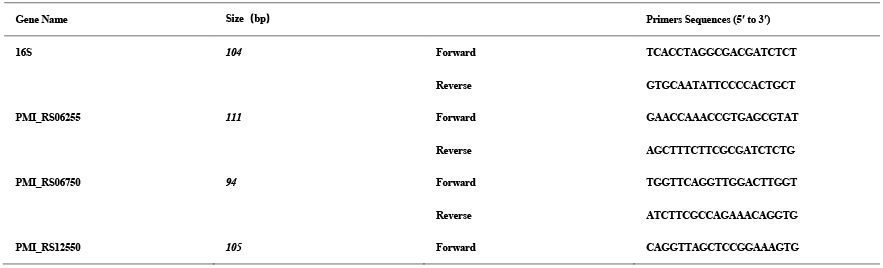
Primers for the target genes used in qRT-PCR

### 2.3 Statistical analysis

The quantitative experiments were performed in triplicates and the data were represented as the mean ± standard deviation (S.D.). A statistical comparison of the data was conducted using two-tailed Student’s t-test. Statistical significance between the two groups was defined as *p* ≤ 0.05.

## 3. Results

### 3.1 Phenotypic characteristics

#### 3.1.1 Electron microscopy findings

Scanning electron microscopy was performed to observe single cell morphology in the PMML and PMGL strains. The results showed that the PMML had no changes intercellular mucus and smooth cell walls than the PMGL strains. (Figure 2).

**Figure 2.**
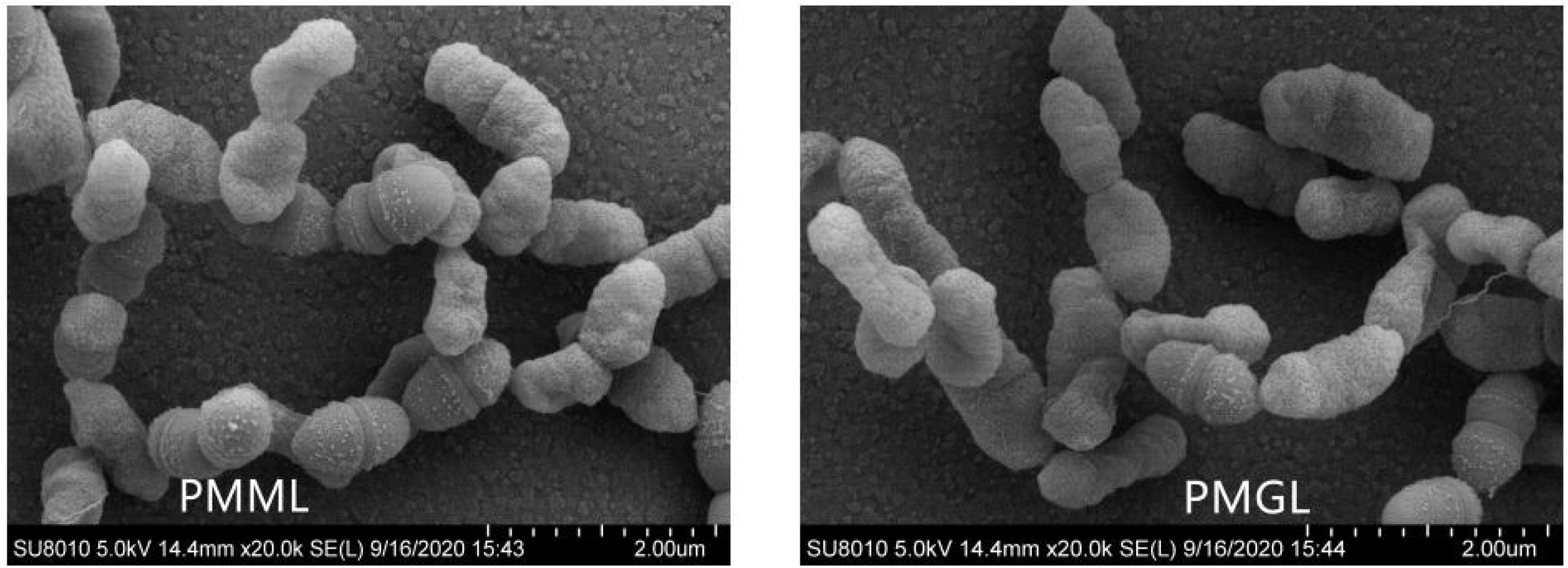
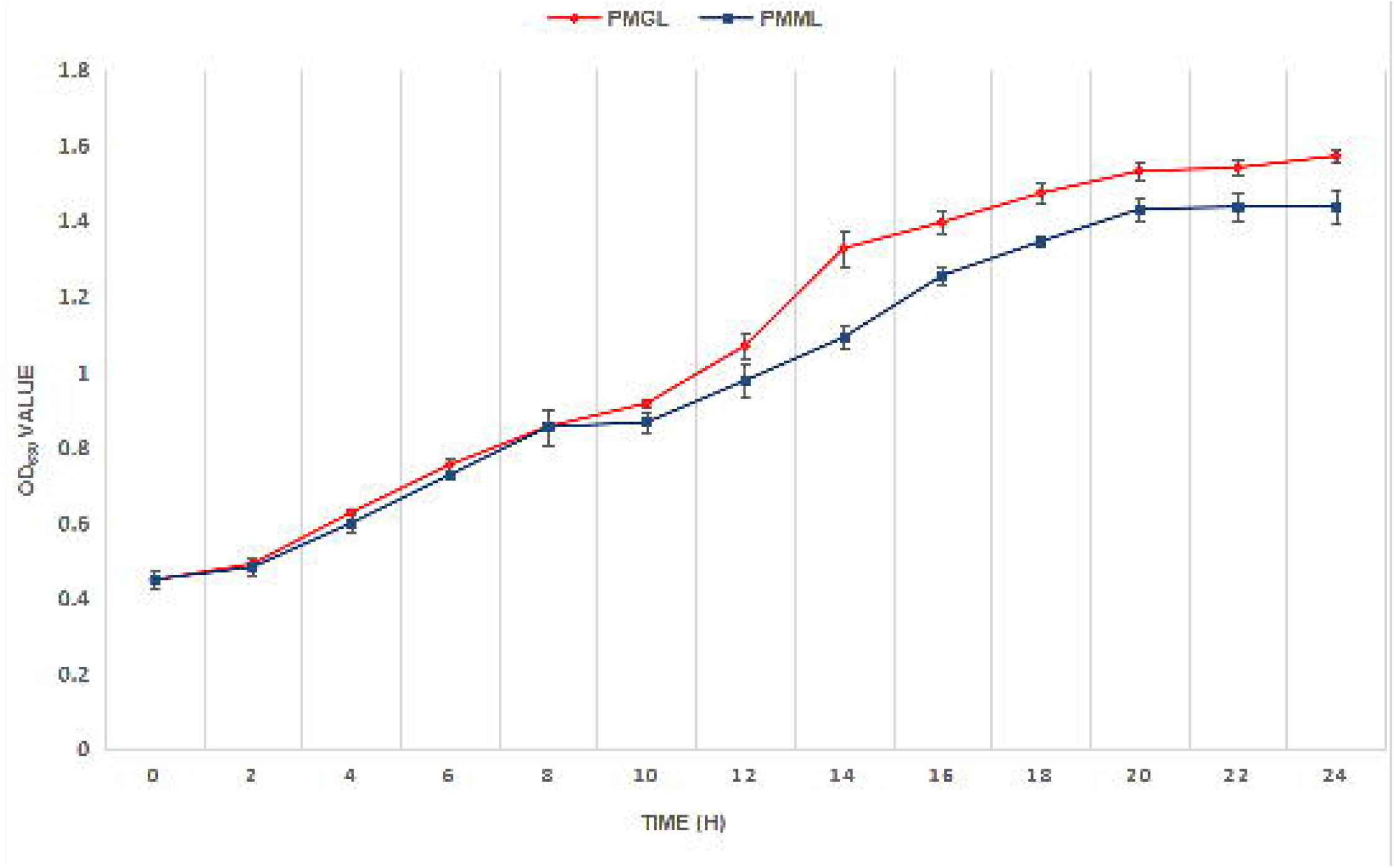
Scanning electron micrographs (SEM) of PMML and PMGL strains. (a) SEM of the PMML strain. (b) SEM of the GS1 strain.

#### 3.1.2 Growth rate assay

A slight difference was observed between the two strains in terms of growth rate with time (Figure 3, Table2). When compared with PMGL, the PMML strain exhibited a decreased growth rate, especially after 14 h (*p* = 0.0095).

**Table 2.** OD_630_ values of PMML and PMGL

| TIME(h) | PMML | PMGL | *p* |
| --- | --- | --- | --- |
| 0 | 0.4497±0.0222 | 0.4480±0.0066 | 0.873 |
| 2 | 0.4823±0.0232 | 0.4913±0.0047 | 0.627 |
| 4 | 0.5987±0.0264 | 0.6270±0.0062 | 0.251 |
| 6 | 0.7270±0.0066 | 0.7537±0.0168 | 0.172 |
| 8 | 0.7780±0.0476 | 0.8553±0.0110 | 0.183 |
| 10 | 0.8663±0.0262 | 0.9167±0.0131 | 0.155 |
| 12 | 0.9770±0.0439 | 1.0697±0.0343 | 0.018 |
| 14 | 1.0917±0.0280 | 1.3280±0.0485 | 0.031 |
| 16 | 1.2560±0.0216 | 1.3973±0.0279 | 0.010 |
| 18 | 1.346±0.0147 | 1.4740±0.0269 | 0.026 |
| 20 | 1.4320±0.0293 | 1.5330±0.0246 | 0.021 |
| 22 | 1.4380±0.0397 | 1.5417±0.0216 | 0.024 |
| 24 | 1.4370±0.0416 | 1.5730±0.0180 | 0.023 |

**Figure 3.**
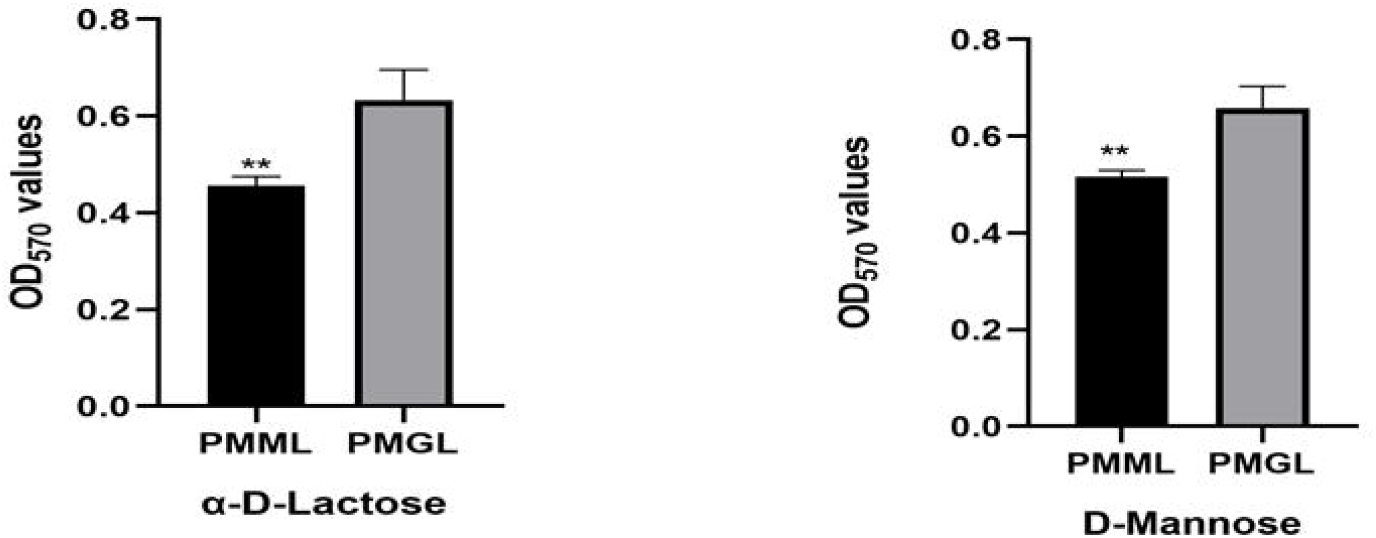
Growth curves of two *P. mirabilis* strains. Growth curves of PMML (blue) and PMGL (red) were determined by measuring the OD_630_ value, which represents the bacterial concentration. The OD_630_ value was measured every 2 h for 24 h.

#### 3.1.3 Chemical sensitivity and carbon source utilization assays

The chemical sensitivity assay showed decreased utilization of α-D-Lactose and D-mannose (*p* = 0.0226 and *p* = 0.046, respectively) by PMML compared to that by PMGL (Figure 4). It is speculated that *P. mirabilis* adapts to a new environment by changing the characteristics of its metabolism, as seen in the simulated microgravity environment. However, the mechanisms involved in this metabolic change are still unclear. Carbon source utilization assays showed that there were no significant differences between other metabolites and stress response pores. However, the Biolog experiment of this study was completed on ground, and not in an actual space environment, and therefore, requires further exploration.

**Figure 4.**
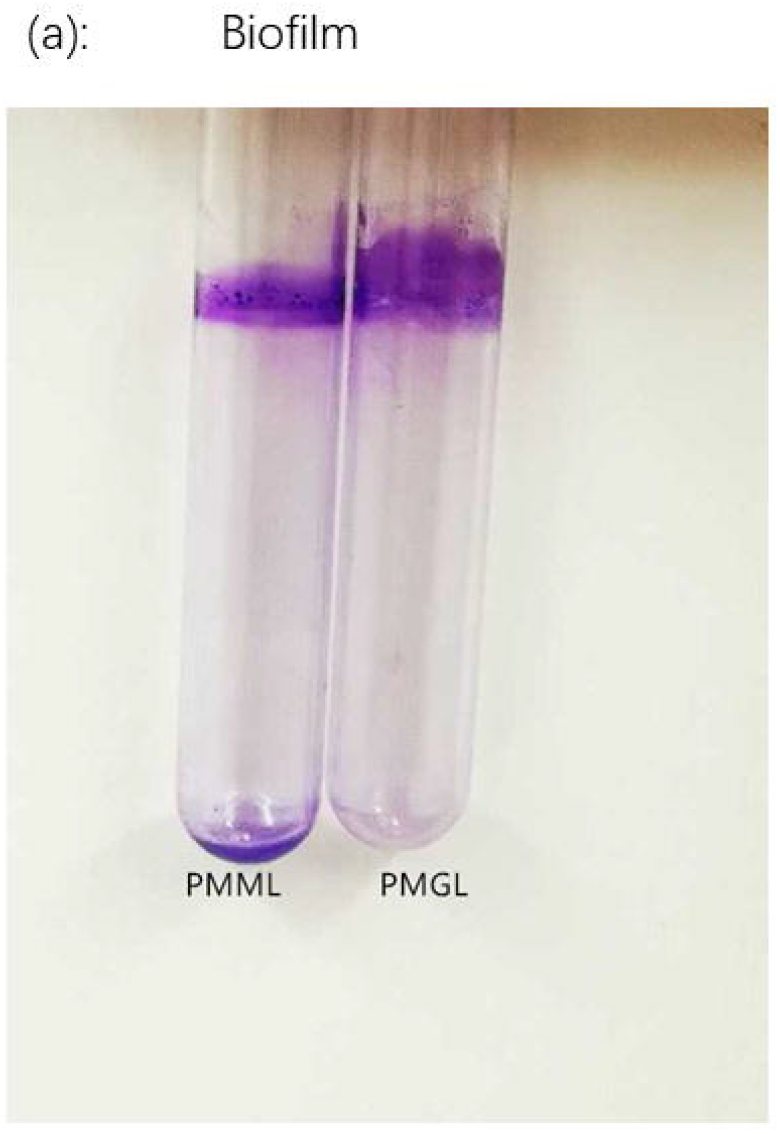
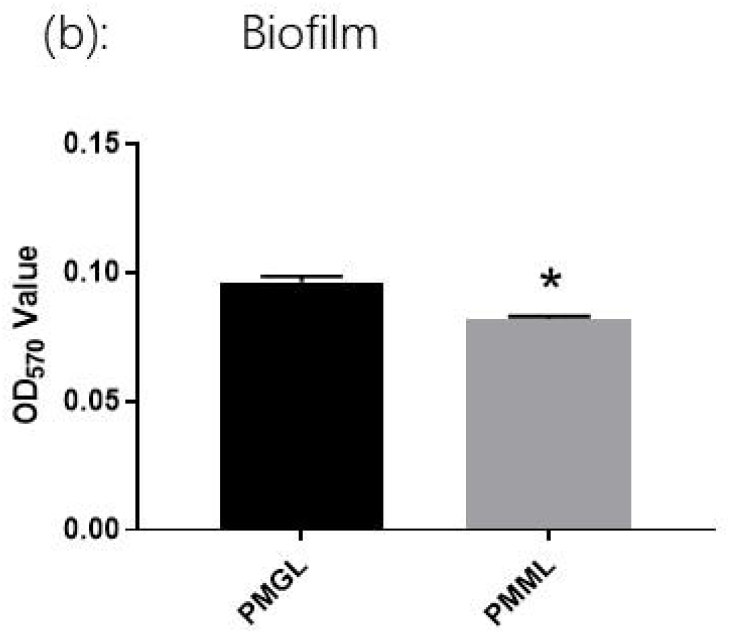
Chemical sensitivity assays of the two strains. The PMML and PMGL strains were incubated in a Biolog GENIII MicroPlate at 37 °C for 24 h in LB medium and tested for carbon source utilization and chemical sensitivity. The effect of simulated microgravity (SMG) on the expression of α-D-Lactose (a) and D-Mannose (b) is shown. Asterisk indicates significant difference at *p* < 0.05.

#### 3.1.4 Biofilm assay

##### 3.1.4.1 The biofilm-forming ability of *P. mirabilis* is considered a significant phenotype

This ability was analyzed in the PMML and PMGL cultures after a 2-week cultivation. The biofilms formed on the glass tubes were monitored by crystal violet staining (a). OD_570_ was measured after crystal violet staining (b). The adhered pellicles were further quantified using the OD_570_ values. PMGL showed a higher biofilm-forming ability than PMML (*p* = 0.0022) (Figure 5)

**Figure 5.**
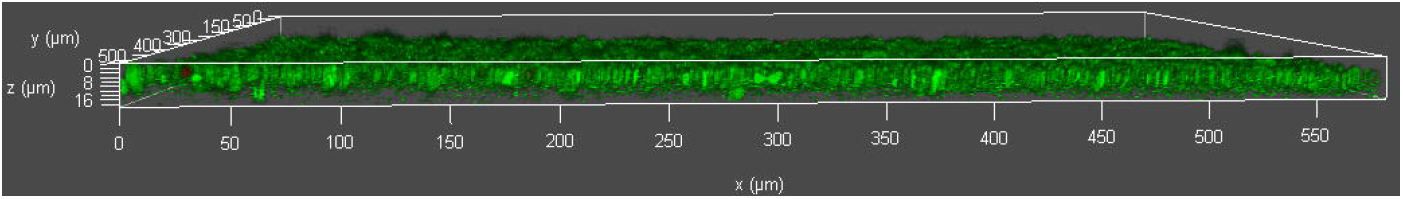
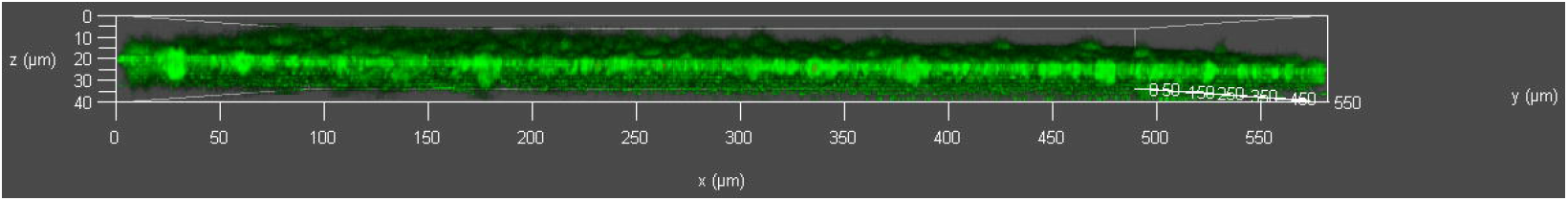
Analysis of biofilm formation ability via crystal violet staining. The PMML and PMGL strains were cultured in 96-well polystyrene microtiter plates at 37 °C for 24 h. Biofilm -forming ability was measured. The biofilms formed on the glass tubes were analyzed using crystal violet staining (a) and quantified by measuring the absorbance of crystal violet at 570 nm (b).

##### 3.1.4.2 Analysis of biofilm formation ability of *P. mirabilis* strain using CLSM

Biofilms of PMML and PMGL strains cultured in 35-mm confocal dishes. Cells were stained with a Filmtracer LIVE/DEAD Biofilm Viability Kit, and biofilm formation ability was tested using confocal laser-scanning microscopy (CLSM). Green fluorescence indicates live cells; red fluorescence indicates dead cells. ***adjusted p value <0.01; *adjusted p value <0.05. (a) Analysis of biofilm formation ability of PMML strain using CLSM. (b) Analysis of biofilm formation ability of PMGL strain using CLSM. (c) Analysis of biofilm formation ability of PMML and PMGL strain using CLSM (Figure 6).

**Figure 6.**
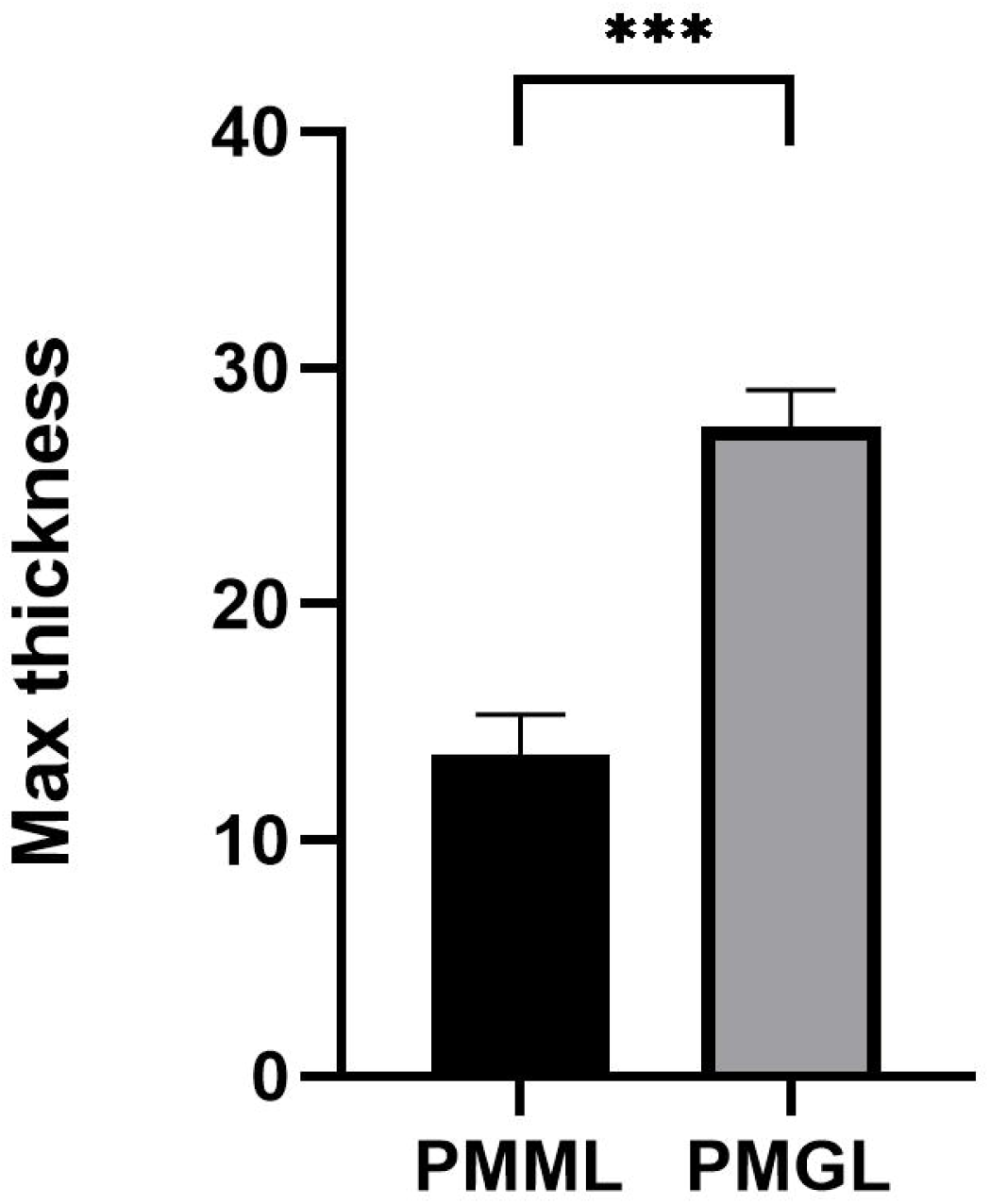
Analysis of biofilm formation ability via CLSM. (a) Analysis of biofilm formation ability of PMML strain using CLSM. (b) Analysis of biofilm formation ability of PMGL strain using CLSM. (c) Analysis of biofilm formation ability of PMML and PMGL strain using CLSM

### 3.2 RNA-Seq mapping and comparative genomic analysis

According to the functional analysis of the KEGG pathway, all DEG clusters were analyzed. Both upregulated and downregulated genes were identified. In PMML, approximately 12 genes were upregulated, and 17 genes were downregulated (Figure 7).

**Figure 7.**
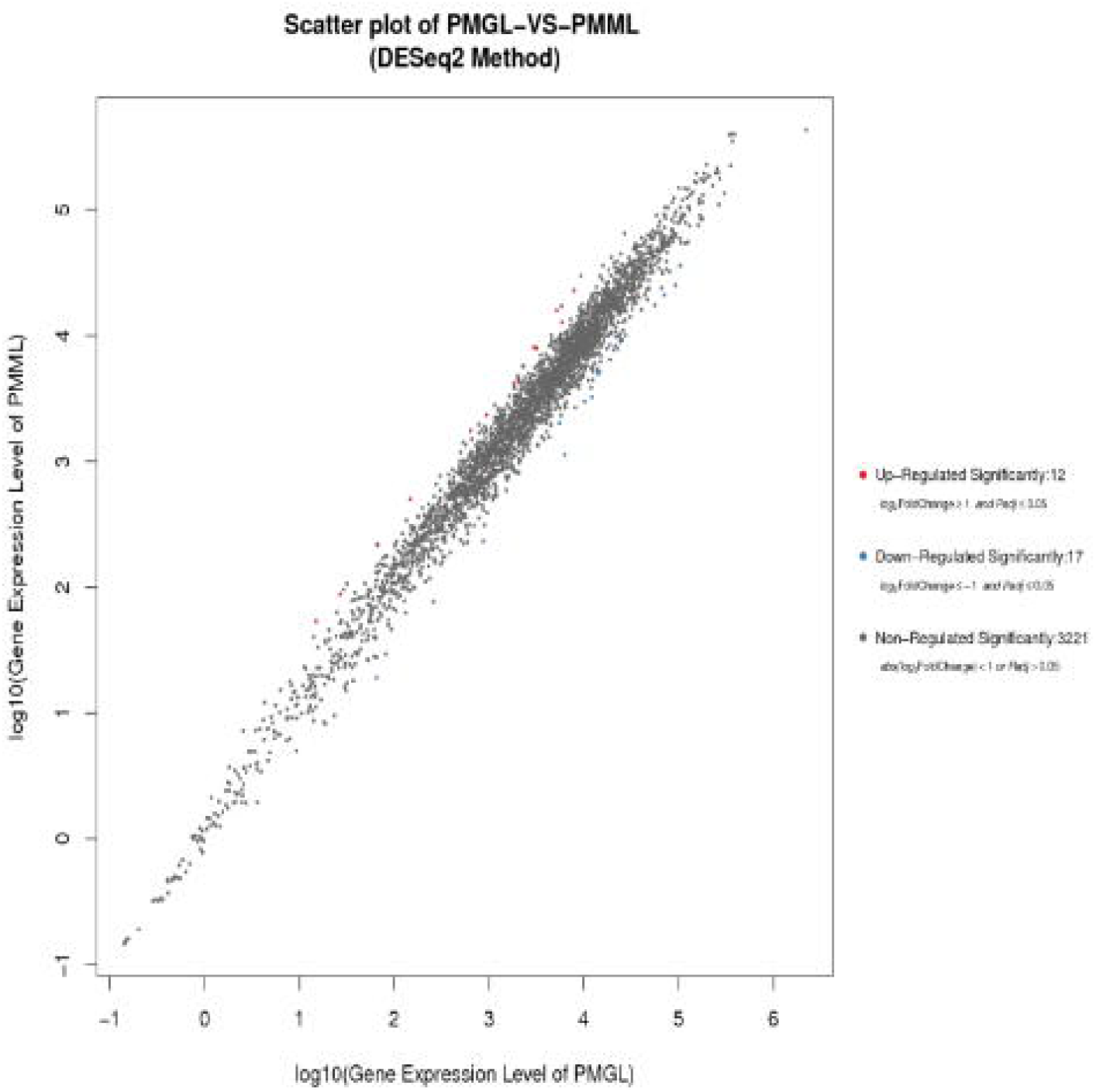
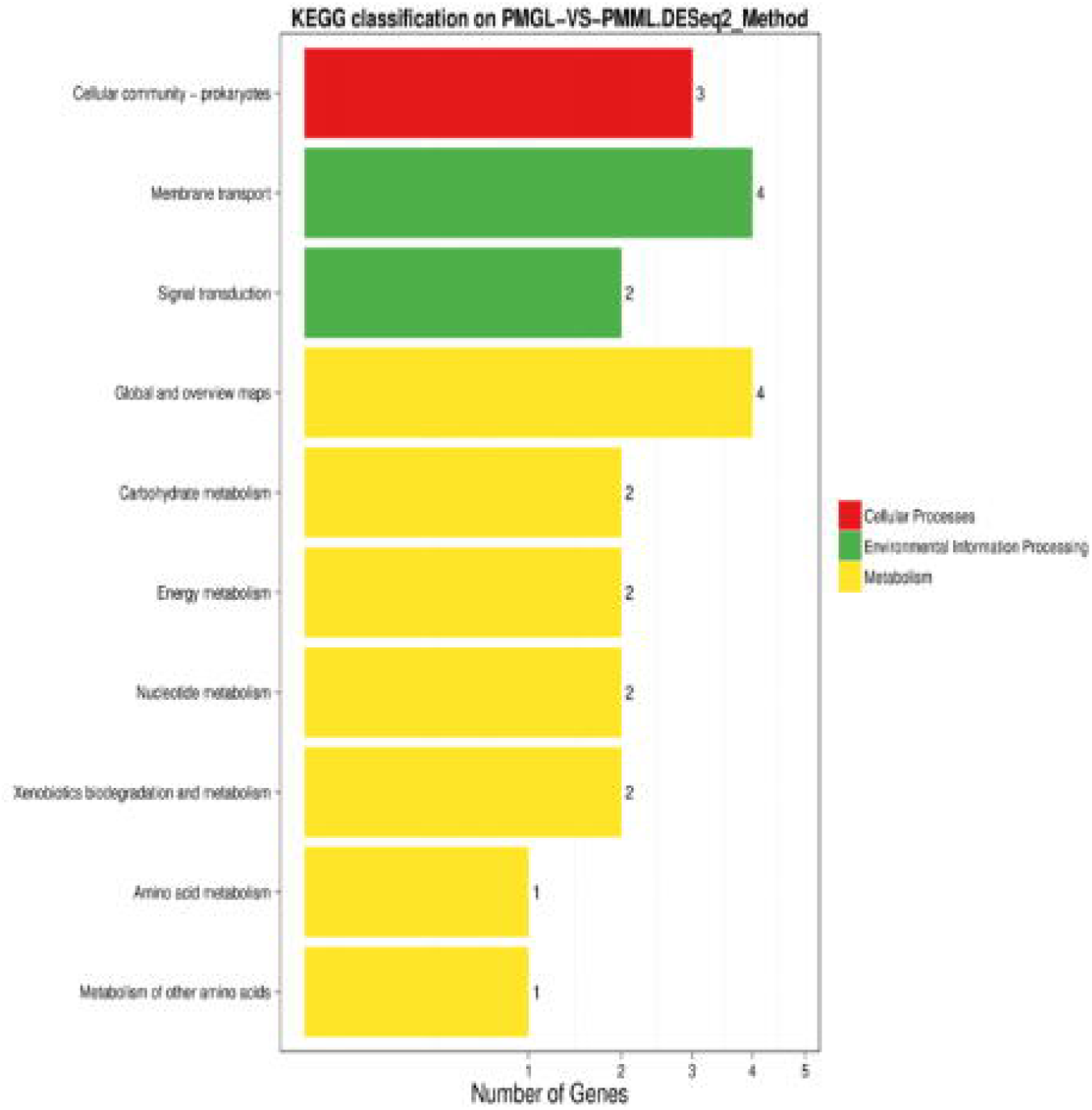
(a) Statistical representation of the differentially expressed genes (DESeq2 Method). The X-axis represents PMML and PMGL; the Y-axis represents number of significant differentially expressed genes. Red represents upregulated genes and blue represent downregulated genes. (b) Global profiling of gene expression changes — Scatter plot of PMGL-VS PMML (DESeq2 Method). The X-axis and Y-axis represents the logarithm of the gene expression level for PMGL and PMML, respectively. Red spots represent upregulated genes, blue spots represent downregulated genes, and gray spots represent the genes that did not change significantly.

We found that the downregulated genes outnumbered the upregulated genes, suggesting that gene metabolism expression was inhibited in PMML. Additionally, most of the DEGs were associated with amino acid metabolism, sugar metabolism, transport, and signal transduction (Figure 8; Table 3). Three DEGs related to biofilm metabolism were identified in PMML, including PMI_RS12550 (*pstS*), PMI_RS06750 (*sodB*), and PMI_RS06255 (*fumC*). The above results were verified using qPCR.

**Table3.**
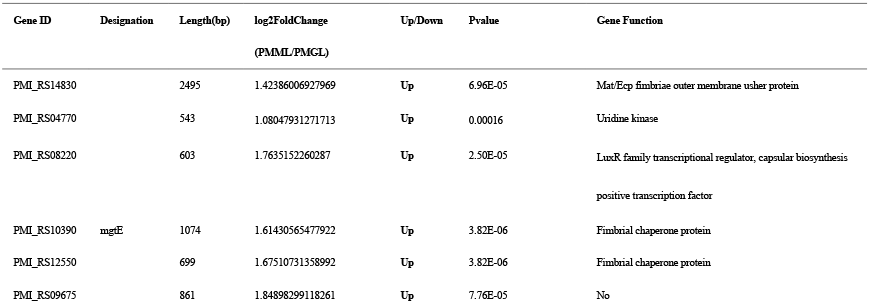
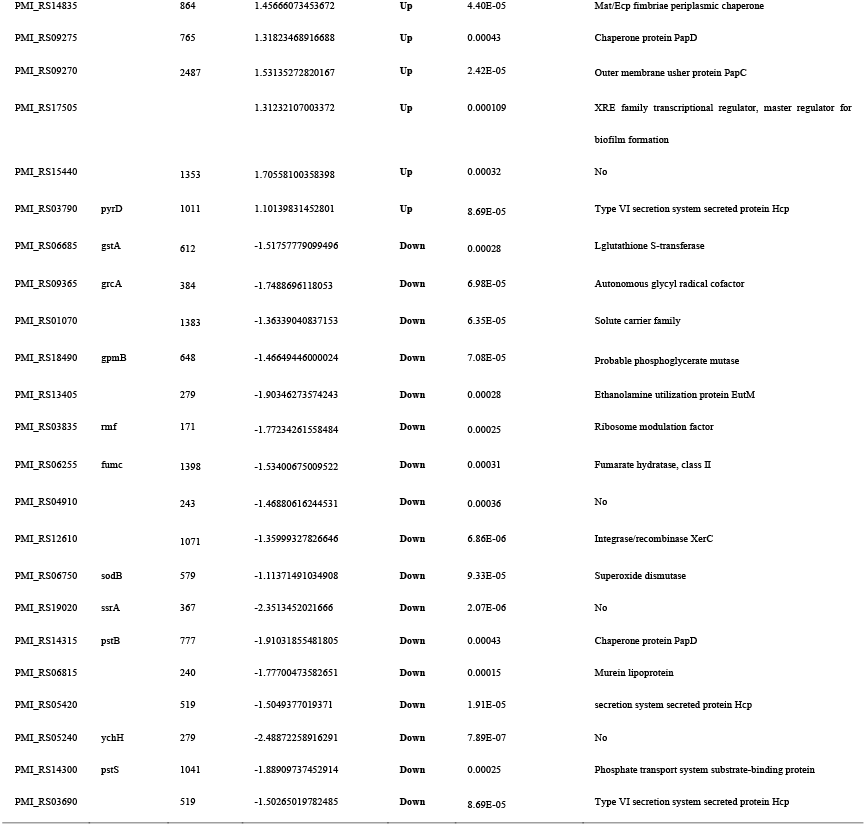
Details of the DEGs.

**Figure 8.**
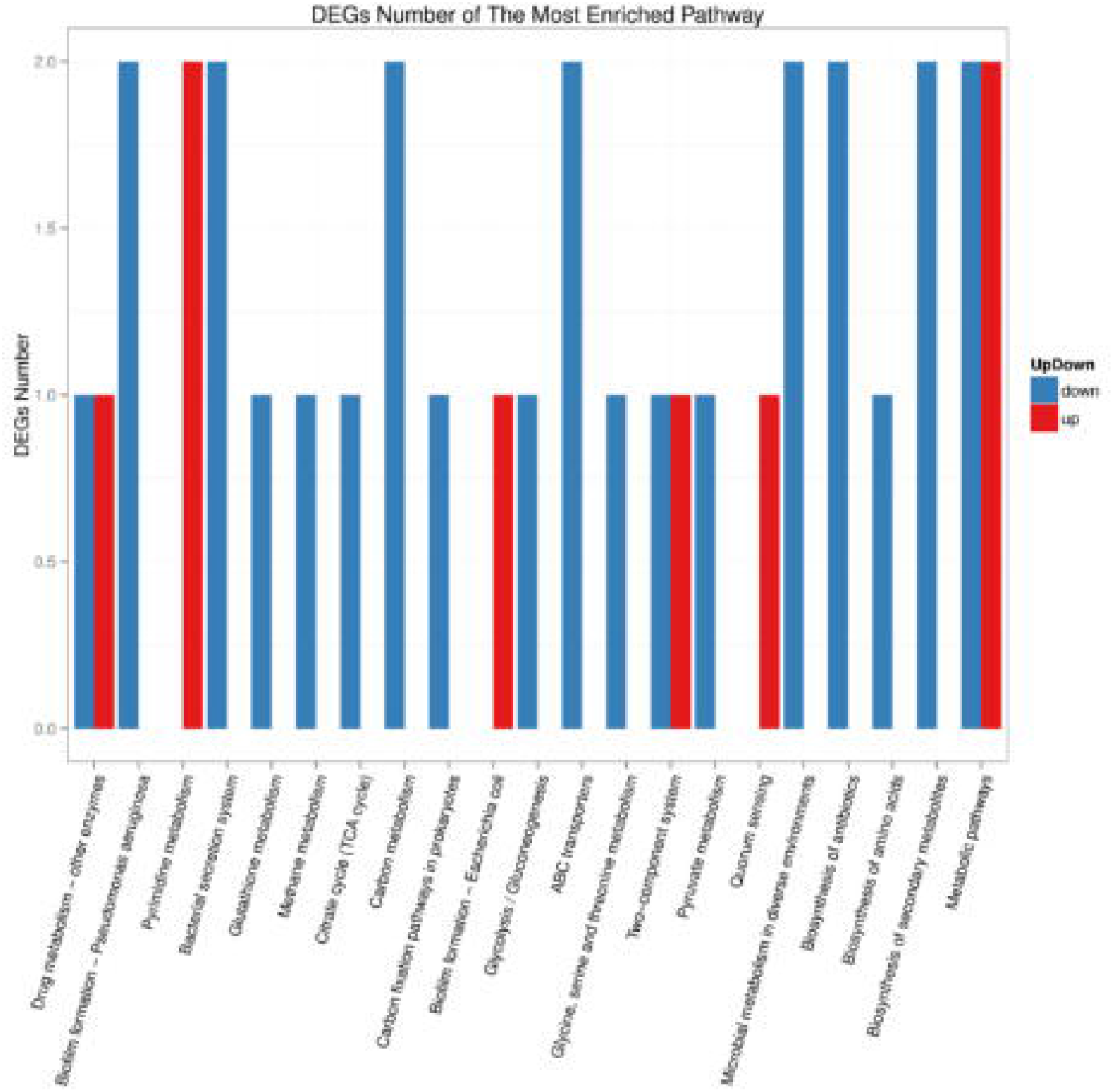
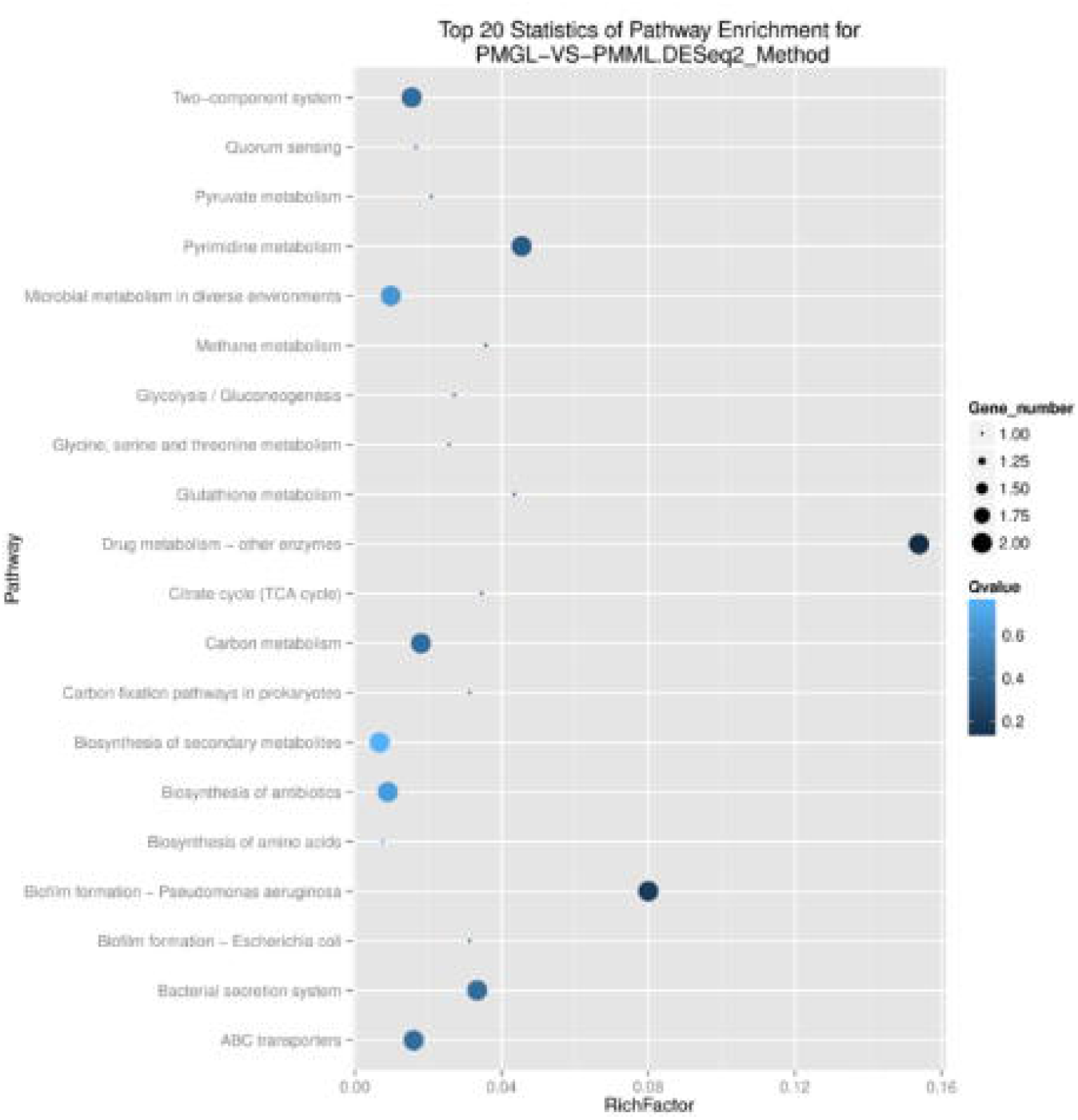
(a) KEGG pathway enrichment analysis of DEGs. The X-axis represents the number of DEGs corresponding to the KEGG pathway. The Y-axis represents the KEGG pathway. (b) KEGG pathway enrichment analysis of upregulated and downregulated DEGs. The X-axis represents the KEGG pathway. The Y-axis represents the number of DEGs corresponding to the KEGG pathways. Blue represents downregulated DEGs and red represents upregulated DEGs. (c) KEGG pathway enrichment analysis of Rich factors. The X-axis represents the number of Rich factors corresponding to the KEGG pathways. The Y-axis represents the KEGG pathways.

### 3.3 mRNA expression levels of key genes related to biofilm formation

The mRNA expression levels of key genes related to biofilm formation were analyzed to determine the mechanisms of biofilm formation by PMML and PMGL. PMML exhibited decreased *pstS, sodB*, and *fumC* levels compared to PMGL (*p* = 0.025, *p* = 0.0056, and *p* = 0.004, respectively) (Figure 9).

**Figure 9.**
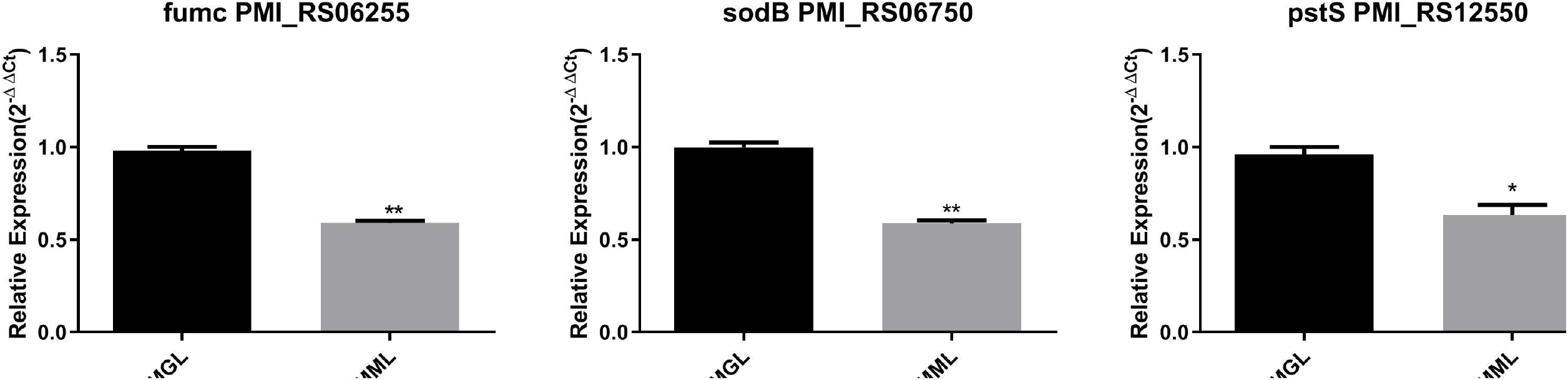
Expression levels of the biofilm-associated genes measured using qPCR. Relative expression is indicated as fold change ± standard derivation between PMML and PMGL. Asterisk indicates the significant difference at *p* < 0.05.

## 4. Discussion

*P. mirabilis* was exposed to simulated microgravity for 14 days using HARV. In addition, its phenotypic characteristics, including morphology, growth rate, metabolism and biofilm formation ability, were analyzed. *P. mirabilis* cultured under the condition of SMG showed a decrease in growth rate, metabolic ability and biofilm formation ability. Further analysis showed that the decrease of biofilm forming ability may be related to the down-regulated expression of genes related to biofilm formation (*pstS,sodB* and *fumC*). It is the first study of *P*.*mirabilis* in SMG environment, providing a certain basis for the prevention and treatment of *P*.*mirabilis* infectious diseases in the future space environment.

Compared with PMGL, the proliferation of PMML was significantly slower than that of PMGL in the logarithmic growth phase. In addition, the growth rate of PMML decreased, which may be related to the down-regulation of the genes involved and changes in metabolism and ribosomal structure. Previous studies have also observed changes in the same rate of growth (Su et al., 2014). In addition, Nicholson et al. (2011) analyzed the growth rate of *Bacillus subtilis* spores on the O/OREOS spacecraft. The flying group of *B. subtilis* cells also showed a lower growth rate compared with the ground control experiment.

The regulation of bacterial metabolism may involve many aspects, including changes in growth rate, utilization of nutrients and changes in metabolites in a complex environment. Studies have shown that α-D-lactose and D-mannose provide necessary energy support in the process of biofilm formation. (Varela et al., 2017, Zhang et al., 2007, Collier and De Miranda,1955. DeGraft-Hanson et al., 1990). In this study, it was found that in the simulated microgravity environment, the utilization rate of two carbon sources (α-D-lactose and D-mannose) decreased. However, the decrease of the utilization rate of α-D-lactose and D-mannose may slow down the biofilm formation or reduce the intensity of biofilm formation. As a result, it further affected the biofilm formation and reduced the growth rate of *P. mirabilis*.

The biofilm formation in bacteria was first carried out under the action of microgravity in 2001 (McLean et al. 2001). Three genes related to biofilm formation, including *pstS,sodB* and *fumC*, were down-regulated in SMG. In theory, *pstS* is a factor related to the ability of bacteria to form biofilm (Cabra et al., 2011). Some people think that the formation of bacterial biofilm may be induced by *sodB* (DePas et al., 2013). Gabryszewski et al confirmed the existence of *fumC* in biofilm and its ability to promote biofilm formation (Gabryszewskietal., 2019). It has been found that *pstS,sodB* and *fumC* may affect the growth rate of bacteria (Rogers et al., 1984, Hassett et al., 1995, Park and Gunsalusso 1995). Studies have shown that the reduction of biofilm formation will affect the viability of bacteria and reduce the growth rate of bacteria (Rogers et al., 1984). A study reports that *pstS* is involved in the formation of biofilms (O’May et al., 2009). The main genes involved in the formation of bacterial biofilm may be *pstS*(Mudrak and Tamayo, 2012; Blus-Kadosh et al., 2013). We believe that due to the stimulation of microgravity, the expression of *pstS* is down-regulated, the biofilm forming ability is decreased, and the biofilm becomes thinner, which may affect the growth rate of *P. mirabilis*. Hassett et al. observed that *sodB* significantly affected the growth rate of *Pseudomonas aeruginosa* (Hassett et al. 1995). *SodB* is a factor related to biofilm formation, and the decrease of *sodB* expression will reduce the biofilm forming ability of bacteria (Chen et al., 2019). In *P. mirabilis*, the environmental changes that inhibit the activity of *sodB* also lead to the weakening of biofilm, which may reduce the growth rate of *P. mirabilis*. Studies have shown that the growth rate of bacteria is affected by the expression of *fumC* gene (Park and Gunsalus,1995). *Fumc* is a kind of II enzyme, which is related to the formation of biofilm (Woods et al., 1988). In our opinions, the decrease of *fumC* expression may weaken the formation of biofilm, resulting in a decrease in the growth rate of *P*.*mirabilis*. As a result, it is speculated that the decrease in the growth rate of *P. mirabilis* under SMG may be attributed to the down-regulation of genes related to biofilm formation.

The growth of bacteria on the surface of microorganisms and the ability of biofilm formation under different growth conditions are affected by many factors, including the type of culture, cutting methods and nutrient utilization. As a result, it showed that *Pseudomonas aeruginosa* showed increased biofilm formation and new structure formation during space flight (Kim et al., 2013). In addition, studies have shown that space flight reduces the biofilm forming ability of multidrug-resistant *Acinetobacter baumannii*(Zhao et al., 2019). Studies have shown that the uncertainty in the treatment of *P. mirabilis* is closely related to the intensity of biofilm formation (Deva et al., 2013). Crystal violet staining and confocal scanning electron microscopy were used to detect the biofilm forming ability of *P. mirabilis*. Compared with PMGL, the biofilm forming ability of PMML decreased. For further analysis at the transcriptional level, a series of the biofilm forming related genes were detected in SMG group. As a result, the decrease of biofilm formation ability of *P. mirabilis* in SMG environment could be attributed to the down-regulation of the expression of the above genes.

## Conclusion

The study provided data showing a decline in growth rate, metabolism and biofilm formation of *P. mirabilis* after 14 days of exposure to simulated microgravity. In addition, further analysis showed that the decrease of growth rate, metabolic ability and biofilm forming ability may be due to the down-regulation of biofilm formation and synthesis (*pstS,sodB* and *fumC*) gene expression. The SMG condition enables us to explore the potential relationship between bacterial phenotype and molecular biology, thus opening up a suitable and constructive method for medical fields that have not been explored before. It provides a certain strategy for the treatment of *P. mirabilis* infectious diseases in space environment by exploring the microgravity of *P. mirabilis*.

## ACKNOWLEDGMENTS

This work was supported by the Key Program of Logistics Research (BWS17J030).

## AUTHOR CONTRIBUTIONS

Dapeng Wang and Po Bai performed the experiments, analyzed the data, prepared figures and/or tables, and authored or reviewed drafts of the manuscript. Bin Zhang and Xiaolei Su performed the experiments and contributed reagents/materials/analysis tools. Xuege Jiang, Tingzheng Fang and Li Xu performed the experiments and prepared figures and/or tables. Junfeng Wang conceived and designed the experiments, authored or reviewed drafts of the manuscript, and approved the final manuscript draft. Changting Liu conceived and designed the experiments, authored or reviewed drafts of the manuscript, and approved the final manuscript draft.

## ETHICS STATEMENT

The study was approved by the insititutional review board (CWO) of the Chinese PLA General Hospital, Beijing, China. Ethical number: S2019-327-01.

